# Multi-continental detection of *Streptococcus pyogenes* M1_UK_: Impact of ssrA SNP on SpeA expression in ancestral and M1_UK_ isolates

**DOI:** 10.1101/2023.04.30.538858

**Authors:** Ana Vieira, Ho Kwong Li, Xiangyun Zhi, Lucy Reeves, Kristin K. Huse, Kai Yi Mok, Oscar Cowen, Elita Jauneikaite, Juliana Coelho, Shiranee Sriskandan, Valerie WC Soo

## Abstract

The *Streptococcus pyogenes* M1_UK_ lineage, characterised by an intrinsic ability to express SpeA toxin and defined by 27 single nucleotide polymorphisms in the core genome, dominates the population of *emm*1 *S. pyogenes* isolates throughout Europe, Canada, South America, Japan and Oceania. While not the sole deterministic factor, enhanced SpeA expression is likely to have contributed to M1_UK_ lineage expansion, but was not sufficient to support expansion of intermediate lineage M1_23SNP_, that expresses SpeA at a similar level to M1_UK_.

A single nucleotide polymorphism (SNP) in the *ssrA* leader sequence upstream of *speA* is one of a limited number of SNPs that distinguish intermediate sublineages that differ in SpeA production. It was recently shown that this SNP leads to increased *ssrA* terminator read-through, and consequent increased transcription of *speA*, which lies downstream of *ssrA*. In this work, introduction of the *ssrA* SNP into representative isolates of the widely disseminated M1_global_ clone and the intermediate M1_13SNP_ lineage, that cannot otherwise produce readily-detectable SpeA in culture, resulted in SpeA expression, confirming the importance of the *ssrA* SNP to SpeA phenotype. Consistent with this, correction of the *ssrA* SNP in M1_UK_ abrogated SpeA expression. However, RNAseq analysis of 8 *emm*1 clinical pharyngitis strains showed that presence of the SNP was not invariably linked to read-through from the ssrA leader sequence or SpeA expression. Read-through was observed in isolates that did not possess the ssrA SNP. Critical review of existing data suggests that *speA* mRNA transcript length may be impacted by the two-component regulator CovRS, pointing to a complex regulatory network interaction between the bacterial chromosome and phage-encoded superantigens.

## Introduction

A marked increase in serious *Streptococcus pyogenes* infections was reported in several countries following relaxation of public health interventions designed to limit the spread of COVID-19 [1]. In England, notifications of both scarlet fever and invasive *S. pyogenes* infections increased throughout 2022; infections peaked at the end of the calendar year but remained high thereafter [2]. In the 30 weeks from mid September 2022-mid April 2023 there were 54,394 notifications of scarlet fever and 2965 invasive *S. pyogenes* infections in England alone [2]. The high case fatality rate of 14.2% [2] underlines the major public health impact of the observed increase, and the importance of understanding the relative roles of strain pathogenicity, population immune susceptibility, co-infections, season, and access to healthcare.

*Emm*1 *S. pyogenes* strains are known to be inherently invasive and normally account for one quarter of invasive infections; however, in the 2022-2023 season, *emm*1 isolates accounted for over half of invasive infections, rising to over 60% in children in England [2]. The population of *emm*1 *S. pyogenes* isolates in England is dominated by a novel sublineage, M1_UK,_ that has an intrinsic ability to express the phage-encoded superantigenic toxin, streptococcal pyrogenic exotoxin (Spe) A [3,4]. M1_UK_ strains can be differentiated from strains belonging to the highly successful *emm*1 *S. pyogenes* globally-disseminated clone that emerged in the 1980’s (M1_global_) by just 27 single nucleotide polymorphisms in the core genome [3]. The M1_UK_ lineage has been identified elsewhere in Europe [5], North America [6,7], Japan [8], China, Russia [9] and Australia, where isolates from Queensland and Victoria date from 2013 [10], just 3 years after its first detection in England [3]. While M1_UK_ represented ∼60% of Australian *emm*1 isolates by 2020 [10], in England, this proportion reached 91% by 2020 [4]. The detection of M1_UK_ Queensland, underlines the ability of this lineage to spread in temperate and more tropical climates, although notably *emm*1 strains appear to be uncommon in sub-Saharan Africa [11]. Of added concern are reports of strains infected with a ΦHKU488.vir-like phage, that carries both *speC* and *ssa* superantigen genes in Australia [10]. To date, this phage has not been identified in *emm*1 strains from England, although ∼10% of *emm*1 strains in England do possess a phage encoding *speC* and *spd*.

Previous genomic analysis of *emm*1 *S. pyogenes* strains in England identified two intermediate sublineages in addition to M1_global_ and M1_UK,_ with just 13 and 23 of the 27 SNPs that define M1_UK_ [3], namely M1_13SNP_ and M1_27SNP_ (Figure 1). We recently reported that strains from the M1_13SNP_ intermediate sublineage produce negligible amounts of SpeA, similar to M1_global_ [12]. Strains from the M1_23SNP_ sublineage, in contrast, produce SpeA at an increased level that is indistinguishable from M1_UK_ isolates [12]. Despite increased SpeA production, M1_23SNP_ isolates could not be detected when screening all *emm*1 isolates submitted to the reference laboratory in England in 2020 [4], suggesting an added fitness advantage in M1_UK_ strains beyond production of SpeA alone. We postulate this further adaptation may be conferred by the additional 4 SNPs that distinguish M1_UK_ from M1_23SNP_ isolates (3 non-synonymous SNPs and an intergenic SNP 39bp upstream of *glpF2*, aquaporin, expression of which is significantly reduced in M1_UK_) [10,12]. In this report we seek to further understand the mechanisms that differentiate SpeA-producers from non-producers among *emm*1 *S. pyogenes* strains that carry the phage that encodes *speA*.

**Figure 1.**
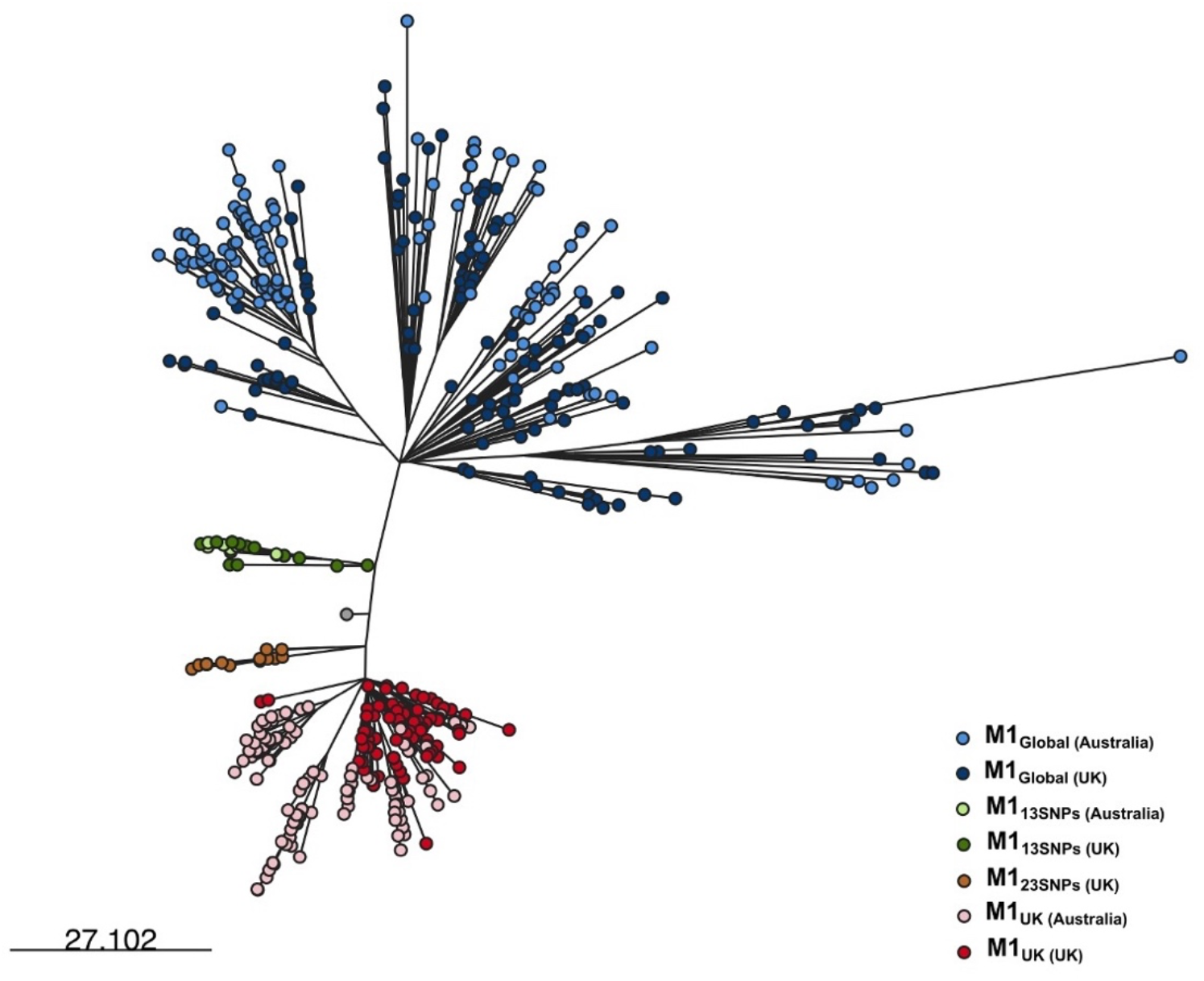
M1_UK_ and sublineages. Phylogenetic tree constructed with 413 core SNPs (without recombination regions) obtained by mapping 269 *emm*1 *S. pyogenes* isolates selected from UK to demonstrate sublineages [12] with 319 isolates from Australia [10] to the reference genome MGAS5005. Coloured tips indicate the various sublineages, with Australian strains indicated by lighter colours and a UK strain with 19SNPs in grey.

## Results and Discussion

The M1_UK_ SNP at position 983438 of the MGAS5005 reference genome, within the leader sequence of *ssrA* and upstream of the *speA* start site (referred to hereafter as the ‘ssrA SNP’), is present in both M1_23SNP_ and M1_UK_ sublineages, but is absent in M1_global_ and M1_13SNP_ sublineages. Screening of natural mutants arising during outbreaks identified the ssrA SNP to be associated with increased SpeA production [12]. Davies et al [10] showed that introduction of the ssrA SNP resulted in increased expression of SpeA by M1_global_. We introduced the ssrA SNP into an M1_global_ strain, and into an intermediate M1_13SNP_ strain (Figure 2A). Consistent with the findings of Davies et al [10], SpeA production in culture supernatants increased in both transformants (Figure 2B). Enhancement of SpeA was similar in both the M1_global_ and M1_13SNP_ backgrounds, suggesting that the accumulated SNPs in the M1_13SNP_ lineage, which include almost half of the SNPs that characterize M1_UK_, do not appreciably contribute to the SpeA phenotype. We considered if our mutagenesis strategy itself might lead to increased SpeA expression but this was ruled out by use of a control construct in each strain (Figure 2B). In contrast, the enhanced SpeA expression in an M1_UK_ background could be reduced considerably upon correction of the ssrA SNP (Figure 2B).

**Figure 2.**
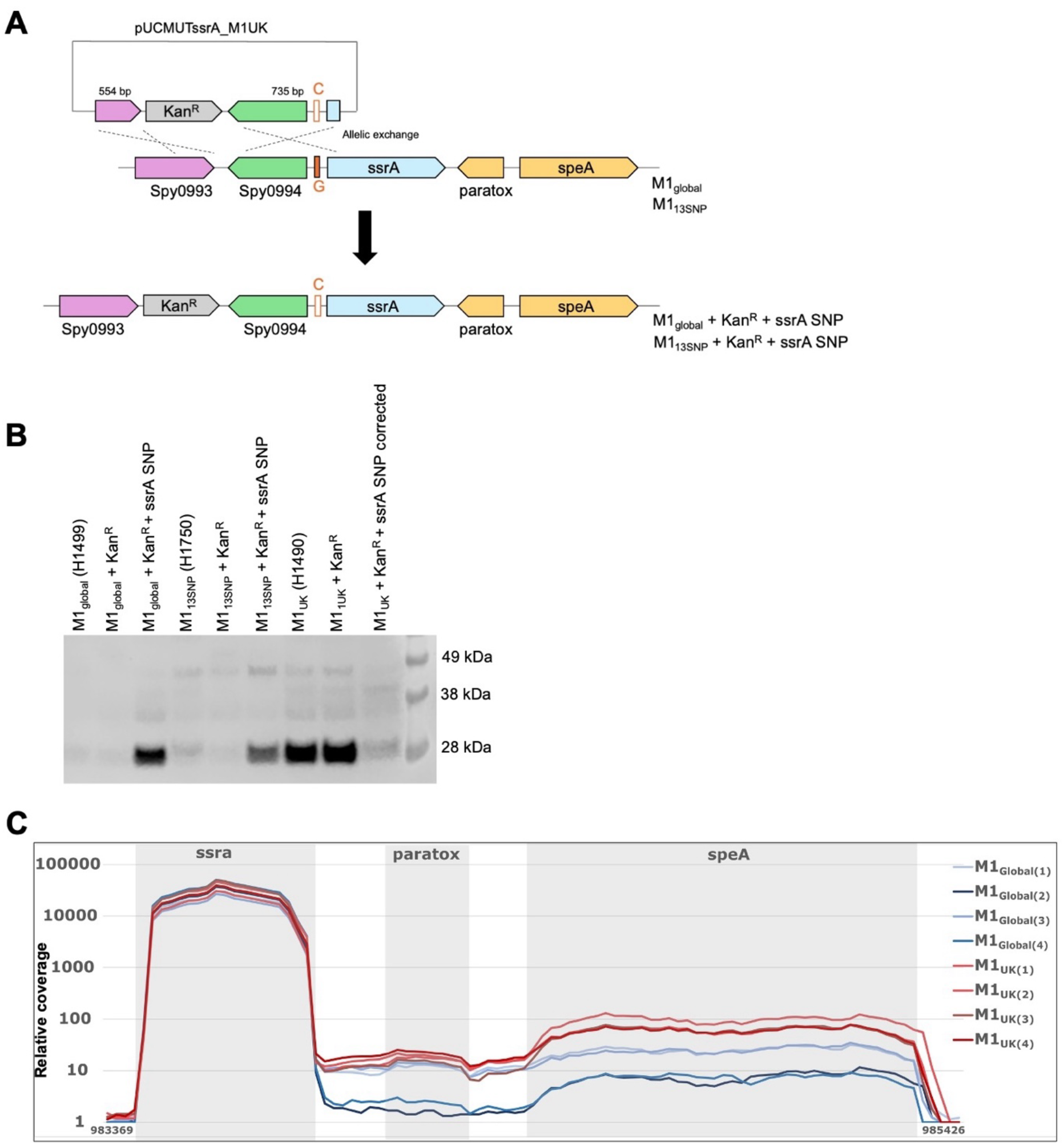
Complex regulation of SpeA production. **A**. The overall schematic for introducing the ssrA SNP into a *S. pyogenes* M1_global_ and M1_13SNP_ lineages. **B**. Introduction of the ssrA SNP into M1_global_ strain H1499 and M1_13SNP_ strain H1750 results in detectable SpeA in supernatants when tested using SpeA immunoblot. AphA3 (KanR) was introduced upstream of the ssrA leader sequence as a selectable marker in all transformants. Only transformants with the ssrA SNP introduced expressed detectable SpeA. In contrast, M1UK strain H1490 expressed considerably lower SpeA upon correction of its ssrA SNP. **C**. RNA sequencing read abundance across *ssrA*; an intergenic region which includes *ptx (paratox*); and *speA* comparing four M1_global_ strains (blue lines) and four M1_UK_ strains (red lines). All were non-invasive throat isolates. M1_global_ isolates were BHS0674 (H1499); BHS0162 (H1489); BHS0130 (H1504); and BHS0151 (H1488). M1_UK_ isolates were BHS0170 (H1490); BHS0128 (H1503); BHS0258 (H1491); and BHS0581 (H1496).

To understand the basis for difference in *speA* expression between the lineages, we compared RNAseq reads from four randomly selected M1_global_ strains, which do not have the ssrA SNP, and four randomly selected M1_UK_ strains, which all have this SNP. None of the strains possessed regulatory gene mutations other than those that characterize M1_UK_. Similar to Davies et al [10], we found that there was indeed evidence of read-through in the intergenic regions between the end of *ssrA* and the start of *speA*. (Figure 2C). However, we did not see clear differentiation between the four M1_global_ strains and the four M1_UK_ strains within the intergenic region, where read abundance in two of the M1_global_ strains was almost identical to M1_UK_ strains (Figure 2C). Regardless, transcription of *speA* was significantly reduced in all four M1_global_ strains compared with M1_UK_. Interestingly, *speA* read abundance was somewhat higher in the two M1_global_ strains that had more intergenic reads, but not as high as the M1_UK_ strains. These data provide support for intergenic read-through affecting *speA* transcription. However, based on short read RNA sequencing data, we cannot definitively ascribe read abundance in the intergenic region to the ssrA SNP in all cases, noting that all four of the M1_global_ strains studied lack any SNP in this region. Comparative WGS analysis between the four M1_global_ strains failed to detect any other genetic features associated with higher levels of read-through. Taken together, transcriptional read-through was observed in this region, but the association with the ssrA SNP and SpeA over-expression was not absolute. Despite not possessing the ssrA SNP, two of four M1_global_ strains tested showed similar read-through to M1_UK_ strains, yet made less SpeA, pointing to additional regulatory factors that influence SpeA expression. The ssrA SNP however was important, at least in the strains examined.

A polycistronic transcript for speA was originally reported ∼25 years ago by Cleary *et al*, who identified both ∼2kB and ∼900bp *speA* mRNA transcripts in a single *emm*1 *S. pyogenes* strain [13]. In that case, the longer 2kB transcript was associated with colonies that failed to produce SpeA, while the shorter 900bp transcript, corresponding to the approximate expected size for the *speA* transcript, was associated with mucoid *emm*1 colonies that produced abundant SpeA. Based on the date of publication, we infer the *emm*1 strain in that study [13] to be from an M1_global_ background. The within-strain variation in *speA* transcript length coupled with mucoid phenotype points to involvement of the two-component regulator covRS (csrRS) which is now known to repress capsule and *speA* transcription in *S. pyogenes* [14–16]. Although M1_global_ strains produce little detectable SpeA protein, SpeA production can be markedly upregulated following mutation of *covS* [14,15] or *covR* [15,16] (Figure 3A), likely accounting for previous reports of upregulation of SpeA following *in vivo* passage of *emm*1 *S. pyogenes* isolates [17].

**Figure 3.**
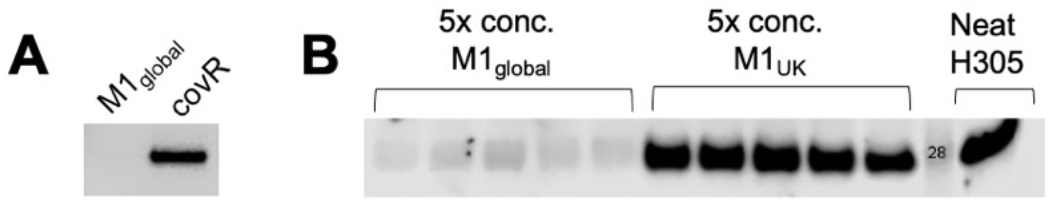
Regulation of SpeA production beyond ssrA SNP. **A**. SpeA immunoblot of supernatant from M1_global_ strain H584 and an isogenic covR T65P mutant (H1565). **B**. SpeA immunoblot of concentrated *emm*1 supernatants from M1_global_ strains (H1488; H1489; H1492; H1495; H1499) and M1_UK_ strains (H1490; H1491; H1493; H1539; H1496) (all non-invasive throat strains) compared with non-concentrated supernatant from *emm*1 strain H305 (NCTC8198).

Consistent with the above experimental observations, contemporary M1_global_ invasive strains with *covRS* mutations arising in patients produce SpeA at amounts equivalent to M1_UK_ strains [12]. A historic *emm*1 isolate NCTC8198 (SF130; H250; H305) [18,19] produces 5-10 fold more SpeA than M1_UK_ strains (Figure 3B) yet does not have the M1_UK_ ssrA SNP. It does however possess a stop mutation in *covS* at residue 318. Northern blotting has previously identified only a single dominant ∼750bp *speA* mRNA transcript for this strain [20], potentially consistent with abundant SpeA production. We postulate that release from indirect or direct covRS repression may allow *speA* transcription to start at a native promoter site proximal to *speA*, triggered by growth-phase related factors. Alternatively, this may result in more rapid processing of any longer transcript. Whether there is an interaction between the ssrA SNP and covRS, other signals, or RNAse Y is unknown.

M1_UK_ is associated with the recent upsurge in invasive infections in Europe [21,22]. The two intermediate sublineages, M1_13SNP_ and M1_23SNP,_ appear to be no longer present in England [4]. Despite the M1_23SNP_ sublineage being able to produce SpeA at a level that is indistinguishable from M1_UK_ isolates [12], it has been fully outcompeted by M1_UK_ in England [4]. Although increased SpeA production is predicted to perpetuate community transmission, the fitness advantage(s) of M1_UK_ may yet require additional features that promote *emm*1 survival in the human population, underlining a need for further research and surveillance.

## Methods and Materials

### Bacterial strains and growth conditions

All bacterial strains used in this study are described in Table 1. *S. pyogenes* isolates were routinely cultured in Todd Hewitt Broth (THB; Oxoid), on THY (THB supplemented with 1% (w/v) yeast extract) agar or Columbia blood agar at 37ºC in 5% CO_2_. *Escherichia coli* One Shot™ TOP10 was grown in lysogeny broth (LB) or LB agar at 37ºC. For maintaining and selecting pUCMUT-containing *E. coli* and *S. pyogenes*, 50 µg/mL and 400 µg/mL of kanamycin were added to the growth medium, respectively.

**Table 1.** Bacterial strains used in this study.

| Strain | Description | Reference/Source |
| --- | --- | --- |
| *S. pyogenes* M1<sub>global</sub> H1499 | Imperial ENT isolate collection | [3] |
| *S. pyogenes* M1<sub>global</sub> H1499 + Kan<sup>R</sup> | Isogenic strain of H1499 with only the Kan<sup>R</sup> introduced upstream of *ssrA* | This study |
| *S. pyogenes* M1<sub>global</sub> H1499 + Kan<sup>R</sup> + *ssrA* SNP | Isogenic strain of H1499 with both the Kan<sup>R</sup> and *ssrA* SNP (G-to-C) introduced | This study |
| *S. pyogenes* M1<sub>13SNP</sub> H1750 | PHE iGAS collection | [12] |
| *S. pyogenes* M1<sub>13SNP</sub> H1750 + Kan<sup>R</sup> | Isogenic strain of H1750 with only the Kan<sup>R</sup> introduced upstream of *ssrA* | This study |
| *S. pyogenes* M1<sub>13SNP</sub> H1750 + Kan<sup>R</sup> + *ssrA* SNP | Isogenic strain of H1750 with both the Kan<sup>R</sup> and *ssrA* SNP (G-to-C) introduced | This study |
| *S. pyogenes* M1<sub>UK</sub> H1490 | Imperial ENT isolate collection | [3] |
| *S. pyogenes* M1<sub>global</sub> H584 | Invasive puerperal sepsis blood isolate | [23] |
| *S. pyogenes* M1<sub>global</sub> *covR* H1565 | Isogenic *CovR* T65P mutant of H584 | [24] |
| *S. pyogenes* M1<sub>global</sub> H1488 | Imperial ENT isolate collection | [3] |
| *S. pyogenes* M1<sub>global</sub> H1489 | Imperial ENT isolate collection | [3] |
| *S. pyogenes* M1<sub>global</sub> H1492 | Imperial ENT isolate<br>collection | [3] |
| *S. pyogenes* M1<sub>global</sub> H1495 | Imperial ENT isolate<br>collection | [3] |
| *S. pyogenes* M1<sub>global</sub> H1504 | Imperial ENT isolate<br>collection | [3] |
| *S. pyogenes* M1<sub>global</sub> H1488 | Imperial ENT isolate<br>collection | [3] |
| *S. pyogenes* M1<sub>UK</sub> H1491 | Imperial ENT isolate<br>collection | [3] |
| *S. pyogenes* M1<sub>UK</sub> H1493 | Imperial ENT isolate<br>collection | [3] |
| *S. pyogenes* M1<sub>UK</sub> H1539 | Imperial ENT isolate<br>collection | [3] |
| *S. pyogenes* M1<sub>UK</sub> H1496 | Imperial ENT isolate<br>collection | [3] |
| *S. pyogenes* M1<sub>UK</sub> H1503 | Imperial ENT isolate<br>collection | [3] |
| *S. pyogenes* M1<sub>global</sub> H305 | NCTC8198 | [19] |
| *E. coli* One Shot™ TOP10 | Cloning host | Invitrogen |

### Introduction of the ssrA G-to-C SNP into the M1_global_ and M1_13SNP_ strains

The ssrA G-to-C SNP was introduced separately into *S. pyogenes* M1_global_ and *S. pyogenes* M1_13SNP_ by allelic exchange (Figure 2A) using the suicide vector pUCMUT [19], which harbours the *aph3* gene (coding for kanamycin resistance cassette (Kan^R^)). *aph3* is flanked by two combinations of restriction sites on either side: PstI + SalI and KpnI + EcoRI.

The 5’ region upstream of ssrA was PCR-amplified from the genomic DNA of *S. pyogenes* M1_UK_ H1490 using primers 993P1 (5’-AA<u>CTGCAG</u>CTATCATCCTTGCATTTGC-3’) and 993P2 (5’-ACGC<u>GTCGAC</u>GGTTTGGTGTTAATTTGATAAAC-3’), and the resulting 554-bp PCR product was cloned into the PstI and SalI sites of pUCMUT, resulting in pUCMUT5s. Next, the 3’ region containing the ssrA SNP was PCR-amplified using primers 994P3 (5’-GG<u>GGTACC</u>TTAATGCTGTAAGAACCAAGATATAA-3’) and 994P4 (5’-AGC<u>GAATTC</u>ATAATGCCTGTCGAATCC-3’), and the 735-bp PCR product was cloned into the KpnI and EcoRI sites of pUCMUT5s to yield the suicide plasmid pUCMUTssrA_M1UK. All constructed plasmids were verified by sequencing at Full Circle Labs.

pUCMUTssrA_M1UK was electroporated into the recipient strains, *S. pyogenes* M1_global_ H1499 and *S. pyogenes* M1_13SNP_ H1750, and was allowed to cross into the chromosome by homologous recombination. The presence of the ssrA G-to-C SNP in the selected strains resulting from a double allelic exchange was verified by PCR and Sanger sequencing (Genewiz).

### Detection of SpeA via immunoblotting

SpeA production in various *S. pyogenes* isolates was assessed using immunoblotting with anti-rSpeA rabbit serum. An overnight culture of *S. pyogenes* (50 mL) was harvested by centrifugation (2,500 rpm, 10 min, 4ºC, swing bucket rotor). The supernatant was passed through an 0.2-µm filter (Sartorius) to obtain a cell-free supernatant. A 500-µL aliquot of this cell-free supernatant was concentrated 5x using a 10 kDa MWCO ultra centrifugal filter (Amicon). The concentrated supernatant was denatured in 1x Bolt™ LDS Sample Buffer (ThermoFisher) containing 300 mM DTT, prior to being fractionated on an 10% Bolt™ Bis-Tris Plus Gel (ThermoFisher). Fractionated proteins were electroblotted onto a nitrocellulose membrane. After blocking the membrane with 5% (w/v) skim milk buffer and 0.05% Tween-20, it was then probed with anti-rSpeA rabbit serum (1:1,000 dilution) [12], and followed by an HRP-conjugated anti-rabbit secondary antibody (1:80,000 dilution). Probed SpeA was detected using chemiluminescence reagents (Cytiva Amersham™ ECL™ Prime Western Blotting Detection Reagents), and the blot was visualised on a Li-COR Odyssey Fc imaging system.

### Quantification of the RNA reads in M1_global_ and M1_UK_ strains

The average read coverage at genomic regions covering *ssrA, paratox*, and *speA* was determined according to Davies et al [10].

## Author contributions

AV, VWCS, HKL, and SS conceived the work described. AV, VS, HKL, XZ, EJ, LR, KKH, KYM, OC, JC (UKHSA) undertook data collection, analysis, and data presentation. AV, VWCS, and SS wrote the first draft of the report and all authors contributed to its revision.

## The authors declare no competing interests

## Acknowledgements

Funding was from UKRI (MR/P022669/1); Wellcome Trust Collaborative award (215539/Z/19/Z); the NIHR Imperial Biomedical Research Centre (BRC). VWCS is supported by an MRC Career Development Award (MR/X007421/1) and HKL was supported by an MRC CMBI Clinical Training Fellowship. The authors acknowledge the support of the NIHR Health Protection Research Unit in Healthcare-associated infections and AMR at Imperial and the Imperial BRC Leonard and Dora Colebrook Laboratory. The authors thank all those who contribute strains to the national and northwest London collections of *S. pyogenes* and the diagnostic and reference laboratories that process these.

## Notes

### Competing Interest Statement

The authors have declared no competing interest.

### Summary of Updates

Title of paper updated to reflect clonal spread to multiple continents Valerie Soo co-corresponding author Methods section included to explain mutagenesis Single multipanel figure now separated into three Mutagenesis figure and western blot updated

